# Foot orientation and trajectory variability in locomotion: effects of real-world terrain

**DOI:** 10.1101/2023.10.18.562999

**Authors:** Emma Gibson, Greg Douglas, Katelyn Jeffries, Julianne Delaurier, Taylor Chestnut, Jesse M. Charlton

## Abstract

Capturing human locomotion in nearly any environment or context is becoming increasingly feasible with wearable sensors, giving access to commonly encountered walking conditions. While important in expanding our understanding of locomotor biomechanics, these more variable environments present challenges to identify changes in data due to person-level factors among the varying environment-level factors. Our study examined foot-specific biomechanics while walking on terrain commonly encountered with the goal of understanding the extent to which these variables change due to terrain. We recruited healthy adults to walk at self-selected speeds on stairs, flat ground, and both shallow and steep sloped terrain. A pair of inertial measurement units were embedded in both shoes to capture foot biomechanics while walking. Foot orientation was calculated using a strapdown procedure and foot trajectory was determined by double integrating the linear acceleration. Stance time, swing time, cadence, sagittal and frontal orientations, stride length and width were extracted as discrete variables. These data were compared within-participant and across terrain conditions. The physical constraints of the stairs resulted in shorter stride lengths, less time spent in swing, toe-first foot contact, and higher variability during stair ascent specifically (*p*<0.05). Stride lengths increased when ascending compared to descending slopes, and the sagittal foot angle at initial contact was greatest in the steep slope descent condition (*p*<0.05). No differences were found between conditions for horizontal foot angle in midstance (p≥0.067). Our results show that walking on slopes creates differential changes in foot biomechanics depending on whether one is descending or ascending, and stairs require different biomechanics and gait timing than slopes or flat ground. This may be an important factor to consider when making comparisons of real-world walking bouts, as greater proportions of one terrain feature in a data set could create bias in the outcomes. Classifying terrain in unsupervised walking datasets would be helpful to avoid comparing metrics from different walking terrain scenarios.

## Introduction

Locomotion is a fundamental human motor pattern that is critical in our daily lives. Control of the lower limbs and feet during locomotion are tightly managed by both the stance and swing limb kinematics and muscle action (1). This process involves both high- and low-level structures of the nervous system, where descending activity integrates with spinal sensorimotor loops that manage the cyclic patterns of the limbs (2, 3). This control allows for adjusting trajectories or orientations of the feet to manipulate the foot-ground clearance, foot placement, foot-ground interaction, and ultimately balance (4-7). This well-controlled but flexible motor program is critical to navigating the ever-changing world around us. When this control fails, a slip, trip, or loss of balance could occur with the potential for ending in a fall (5).

Much of the underlying research investigating foot motion in locomotor contexts was gathered in laboratory conditions with various experimental setups using both treadmill and over-ground walking (8). While important in understanding the neuromechanical control of foot motion while walking, these conditions don’t capture the dynamic and variable environments we normally navigate in our daily life. With the evolving technology of wearable motion capture systems, measuring body kinematics is not only becoming more accessible, but it can be done in real-world contexts (9). By leveraging this technology there is potential to collect large datasets during free-living and unsupervised walking, which could have implications for understanding how clinical conditions or functional changes impact locomotion over larger time scales (10, 11). Untethering these data collections from the laboratory environment gives ready access to more diverse walking scenarios that can differ in terrain and context.

The last decade has seen a growing body of literature evaluating foot biomechanics while walking in real-world, or at least out-of-lab conditions. Frequently, this is achieved by using an inertial measurement unit (IMU) affixed to the foot (12). The acceleration, angular velocity, and in some cases, magnetic field signals from the IMU are combined using various strategies to calculate the orientation and trajectory of the foot (13-15). Validating these algorithms and experimental setups is an important step in development, which has largely occurred in laboratory settings given that the ground-truth comparison is typically an optoelectrical motion capture system. Under controlled conditions of a lab, foot orientation can be estimated with an IMU to within 2.5° depending on the axis of interest (13, 16), while foot trajectories are accurate to approximately 20mm (17). Under more varied or complex conditions that are typical of real-world walking we see slightly inflated error; for example horizontal plane foot angles can exhibit errors up to 3.1° (18) and horizontal trajectory estimates within 1% (19). These small errors in estimation are largely outweighed by the advantage of being able to collect data in vast quantities and in nearly any environment.

A challenge facing real-world, unsupervised gait research is that the features of our built world can influence walking biomechanics to varying degrees (20). Therefore, we need information on how different terrain conditions affect gait, otherwise it becomes difficult to separate changes due to person-specific factors (e.g., disease, injury, functional decline) from simple changes in context or terrain. Initial developments in this area used foot mounted IMUs to classify what terrain the person was walking on as they navigated inclines, stairs, and flat ground (21, 22). Both studies used foot trajectory and gait event timing to generate the classification with accuracies above 95%. It then becomes important to understand the extent in which terrain conditions affect foot biomechanics. Kowalsky, Rebula (23) investigated this during level ground walking while varying the surface features (concrete, gravel, dirt, grass, and woodchips). These authors noted that overall foot trajectory including foot clearance, stride lengths, and variability of these measures tended to increase under more challenging terrain (e.g., gravel vs concrete). While these studies are building the necessary fundamental methods and data to advance real-world gait research, we still lack a direct comparison of foot motion among geometrically different terrain, including flat ground, slopes, and stairs.

The objective of this study was to examine how foot trajectory and orientation change while walking on commonly encountered terrain in daily living activities. This was an exploratory study seeking to inform more directed, hypothesis-driven research in the future. However, we expect the physical constraints of the stair conditions will limit stride length and width compared to the slope and flat ground walking conditions.

## Materials and methods

This study was a cross-sectional, experimental study of the walking mechanics exhibited by healthy adults while navigating typical environments we encounter in our daily lives. We recruited participants on a convenience basis from the surrounding university between November 2021 and august 2022. Prior to any data collection, participants provided written informed consent prior to beginning the methods that follow. The study was approved by the institutional ethics review board (H21-03053).

To be eligible, participants had to be at least 18 years of age, capable of walking intermittently for 60 minutes, and able to fit into the provided shoes which housed the IMUs (sizes from women’s 5 to men’s 13). We excluded any person who reported recent lower body injuries or surgical procedures (previous 12 months), had an acute or chronic condition that inhibited normal physical activity levels or affected gait, or who required a gait aid.

### Instrumentation and equipment

To record foot biomechanics during walking, we instrumented shoes with a wireless IMU embedded in the sole of each shoe, described in previous work (24, 25). The sensor (MPU-9150, InvenSense, CA, USA) measured acceleration (signal range: ±4g), angular velocity (signal range: ±500°/s), and magnetic field strength (signal range: ±1200μT) and a microprocessor (STMicroelectronics, STM3F401, Switzerland; clock rate=84MHz) sampled these signals at 100 Hz before writing them to the onboard microSD chip (MicroSDHC Class 4, Transcend, China). The printed circuit board and components were housed in a custom 3D printed shell with an external power switch, reset button, and charging port. This system was embedded within the sole of both the right and left shoes, where a relief was cut to accommodate the sensor (Fig 1). The relief was positioned to minimize axial loads under the heel and bending forces from the shoe throughout the gait cycle. The sensor y axis (green arrow) was aligned to the long axis of the foot, the x axis (red arrow) pointed to the right on the plane of the shoe’s sole, and the z axis (blue arrow) pointed vertically following the right-hand rule.

**Figure 1.**
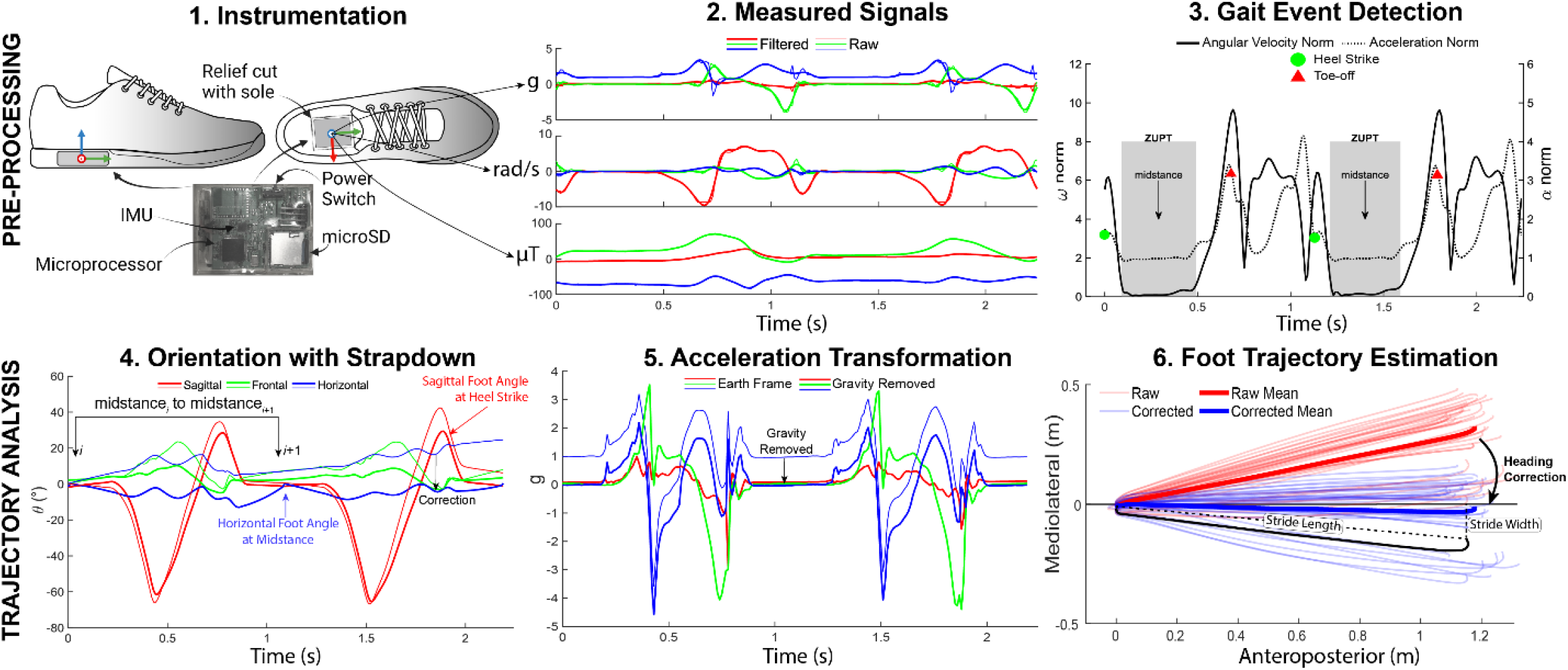
Methods overview. An IMU was fit securely into a relief cut out into the sole of the shoe (Step 1). The forward axis of the shoe (y-axis) was aligned with the forward axis of the foot using previously established methods (24, 26). Each sensor recorded 9 axes of data (Step 2) which were filtered and segmented based on gait events (21, 27). Orientation estimation, shown in Step 4, was performed over each stride using a modified strapdown integration which estimated orientation in the forward and backward directions (28, 29), and then linearly weighted the two signals based on the distance from the start or end of the stride, respectively. Step 5 shows the acceleration signals from Step 2 transformed into the earth frame of reference and with gravity removed (blue lines). Finally, double integration yielded foot position in the mediolateral and anteroposterior directions, with an origin in the local frame (foot position at the previous midstance). These trajectories were corrected for the overall heading based on the global trajectory from the start to the end of the walking bout (assumes linear walking, which was constrained in this experiment). Step 6 shows the results of performing this analysis over all the strides of a participant for one walking condition. The thick lines represent the mean stride trajectory.

### Data collection

While wearing both instrumented shoes, participants performed a series of linear walking bouts on four different terrain conditions (Fig 2). Participants performed each condition, except the stair navigation, at a self-selected, fast, and slow walking speed; however, for the purpose of this study we only analyzed the self-selected speeds due to increased error in gait event detection and orientation at these other speeds. For the slope and stair conditions, both an ascending and descending trial were collected.

**Figure 2.**
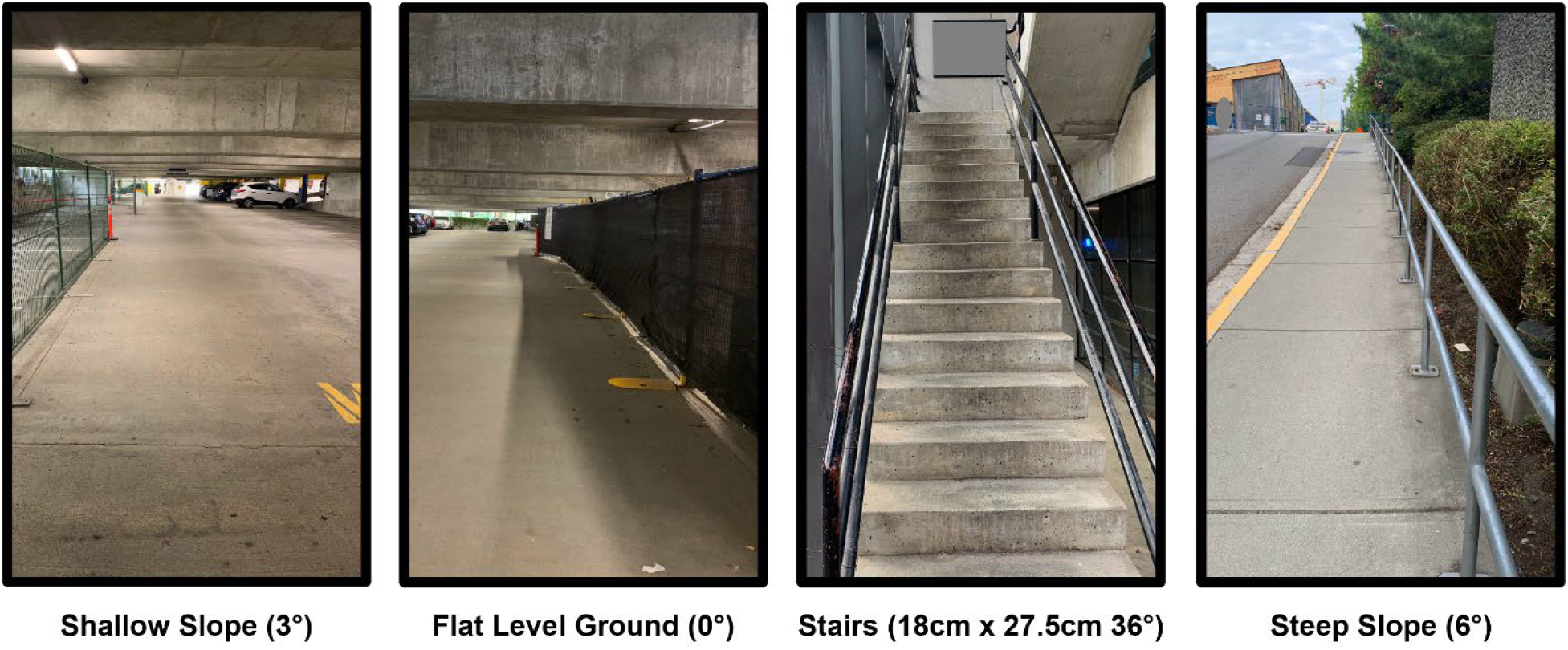
Terrain conditions used in the experiment. A total of seven conditions were collected including ascending and descending for the slope and stairs, and the flat ground condition.

### Data analysis

Drawing on previously published and validated methods, we developed a multi-stage analysis to calculate foot trajectory and orientation estimates of each foot from the raw IMU data. Analyses were performed in MATLAB (v 2021b, Mathworks, Natick, MA, USA). We extracted a series of IMU derived outcomes that are described in Table 1. Each outcome was calculated on a per-stride basis and summarized as the mean or standard deviation to quantify the magnitude and variability for each condition. The following methods were used to process signals, identify gait events, calculate foot orientation, and foot trajectory.

**Table 1.** Definitions of the extracted outcome measures.

| Outcome | Unit | Description |
| --- | --- | --- |
| Stride length | meters | Euclidean distance of the foot position at consecutive midstance events measured in the earth frame local to the current stride (position is zero at start of stride) |
| Stride width | meters | Absolute mediolateral distance of the foot position at consecutive midstance events measured in the earth frame local to the current stride (position is zero at start of stride) |
| Stance time | seconds | Time between heel strike and successive toe-off gait events |
| Swing time | seconds | Time between toe-off and successive heel strike events |
| Cadence | strides/minute | Average strides taken each minute |
| Horizontal plane foot angle at midstance | ° | Differences between the yaw angle of the foot at midstance and the walking heading direction, commonly referred to as foot progression angle |
| Sagittal plane foot angle at HS | ° | Pitch angle of the foot at heel strike, commonly referred to as foot strike angle |

#### Pre-processing

Each walking bout (21 per limb, 42 per participant) was recorded to a single data file with quiet standing for 1-2 seconds at the start and end of each file. Signals from the IMU were filtered with a bidirectional Butterworth lowpass filter (cut off = 10 Hz, order 4) and these filtered signals were stored for use in identifying gait events. The angle between the long axis of the foot (center of heel to center of the toe-box) and the IMU’s y-axis was calculated and used to align the IMU axes to the foot, based on previous methods (24, 26). Our analyses that follow made the assumption that the IMU frame and foot frame were coincident.

The filtered angular velocity about the mediolateral axis of the sensor was then used to identify the positive peaks during the swing phase of locomotion (mid-swing peak). First, the negative peaks of this signal, characteristic of initial contact and toe-off, were roughly identified and the average time between each negative peak was calculated. The mid-swing peak indices of this signal were then identified based on the maximum positive value that were separated by at least 75% of the average time between negative peaks. This assisted with eliminating erroneous mid-swing indices.

#### Gait event detection

Gait events (initial contact, midstance, and toe-off) were determined using two previously published strategies, one for stairs (21) and the other for the slope and flat ground conditions (27). For stair conditions, the toe-off event was identified first based on the second negative peak in the mediolateral angular velocity signal that occurred between two mid-swing events.

Thereafter, midstance was estimated as the minimum of the angular velocity signal energy using a 200ms sliding window. The terminal swing peak deceleration index was found based on the first minimum of the anteroposterior acceleration signal from mid-swing to midstance. Initial contact was calculated as the peak of the squared anteroposterior deceleration signal between the terminal swing deceleration event and 60% of this event to the midstance event.

Slope and flat ground walking produced a more consistent and characteristic waveform which allows for a more simplistic event detection algorithm. Initial contact was first approximated by the first negative peak of the mediolateral angular velocity signal. A 20-sample window around this point was then searched to locate the minimum value of the acceleration Euclidean norm. Midstance was found using a threshold approach for the angular velocity (1.7 rad/s) and acceleration Euclidean norms (1.1 m/s^2^) (19). Finally, toe-off was found in the same manner as initial contact, but the window was centered around the second negative peak of the mediolateral angular velocity signal. A single stride was the time from a midstance instance to the subsequent midstance.

#### Orientation estimation

The orientation of each foot with respect to the earth frame 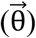 was estimated by sensor fusion using a complementary filter and represented by Euler angles (x, y, z sequence). This approach combined the favorable features of the accelerometer, gyroscope, and magnetometer. The filter worked in three steps: 1) estimate orientation based on acceleration 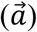 and magnetometer 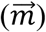 signals 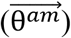), 2)estimate the orientation 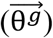 by integrating the angular velocity 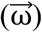 with the sampling time (Δt = 0.01) and adding it to the previous orientation estimate 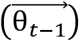, and 3) linearly combine these two orientation estimates with a weighting factor (β = 0.985). Foot orientation was calculated for each sample point, t ∈ [1 … *N*], within a single stride where *N* is the number of data samples between the time points of the *i*^*th*^ midstance and the *i*^*th*^ + 1 midstance.

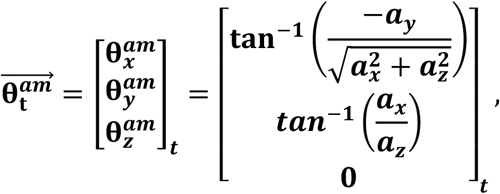

where ***a*** is the normalized acceleration signal in the ***x, y, z*** axes.

Prior to calculating θ_*z*_, the magnetic field vector was transformed into the earth frame based on the currently estimated tilt orientation of the foot.

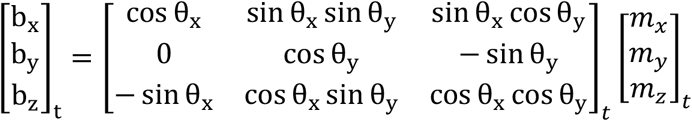

where *m* is the normalized magnetic field strength vector.

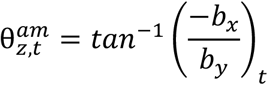

Next, sensor orientation was calculated by integrating the angular velocity signals over each time point and adding it to the previous orientation estimate. For the first time step of a stride, the previous orientation 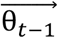was initialized using 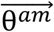.

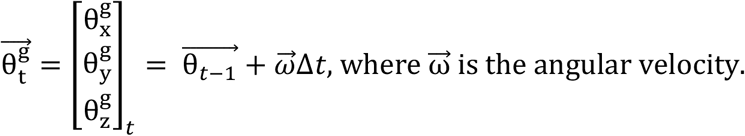

The two orientation estimates in the foot frame were combined to yield the current orientation for time point *t*:

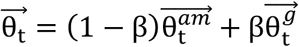

Each orientation estimate was then converted into its quaternion representation (*q*_*t*_) using scalar first notation. This was done to simplify implementing the next processing step, a modified strapdown integration.

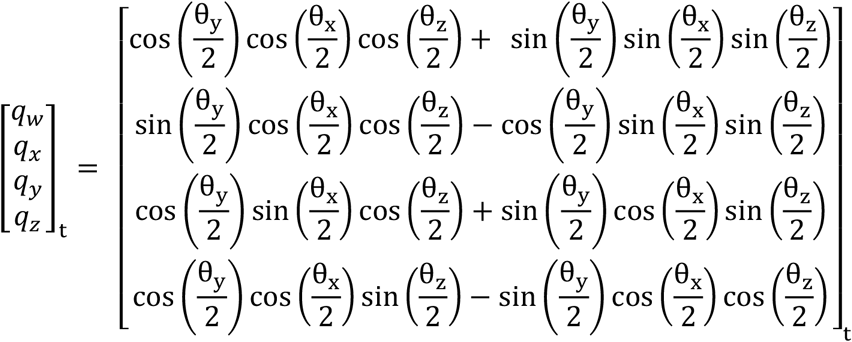

The orientation estimation was augmented by a modified strapdown integration method described by Falbriard, Meyer (28). The orientation estimation outlined above was performed in both the forward and backward directions with respect to time, for each stride independently. Then a linear combination of the orientation estimates arising from the forward (*q*^*f*^) and backward (*q*^*b*^) directions was used to correct the orientation based on the nearest time point with a zero-velocity update point (*i* or *i*+1 midstance point).

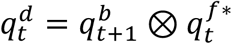, where * is the conjugate and ⊗ is the quaternion product.

The weighting of the forward and backward orientation estimates was achieved by modifying the helical angle, *h*_*t*_, of 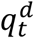. More weight was given to the forward integration for time points nearer the beginning of the stride, while the opposite occurred at time points nearer the end of the stride.

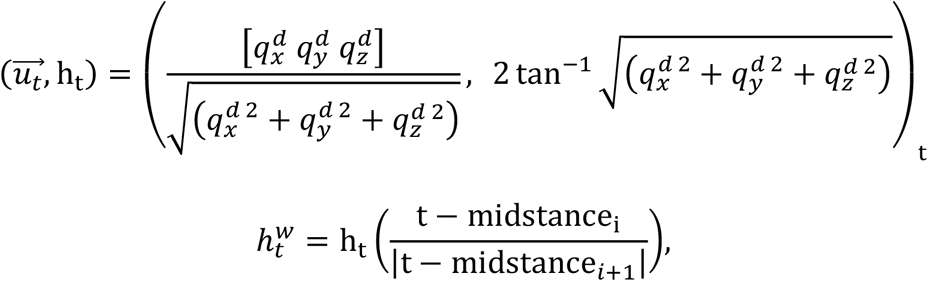

where 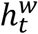 is the weighted helical angle that is transformed back into quaternion form and multiplied with the forward-estimated orientation (*q*^*f*^) to yielf the final orientation of the foot with respect to the earth reference frame (*q*):

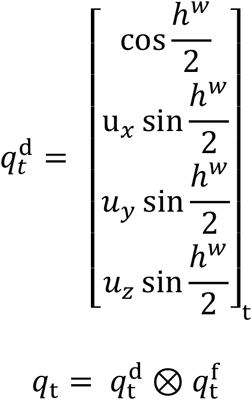

This elicited a corrected quaternion for each sample (*q*_*t*_). For the purposes of reporting foot orientation, we extracted Euler angles in the x, y, z order, where the angles about each axis represent sagittal, frontal, and transverse rotations, respectively. This process was performed on each stride across all walking conditions and for both right foot and left foot IMUs.

#### Gravity correction and trajectory estimation

To determine linear accelerations of the IMU, the raw acceleration data was transformed into the earth reference frame based on the orientation, and the gravity vector was removed to yield linear accelerations in the earth frame:

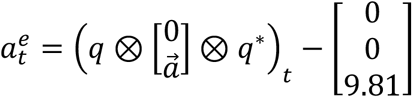

The linear acceleration was integrated over the time steps from the start to the end of the current stride to yield linear velocity. However, these velocities are prone to drift, due to accelerometer bias, which can lead to a non-zero IMU velocity at the next midstance event. Based on a zero-velocity assumption at each midstance, we applied a previously reported correction (29). These corrected velocities were again integrated to yield position.

Finally, the linear real-world heading was set to zero such that the IMU trajectories were centered about the heading (no deviation from the heading equated to zero in the mediolateral trajectory). We achieved this by taking a linear fit of the mediolateral and anteroposterior trajectory data at each midstance event over the duration of the walking bout and then removed the interpolated linear fit from each trajectory datapoints. The resulting foot trajectories (representative participant shown in Fig 3) were used to calculate discrete variables on a stride-by-stride basis.

**Figure 3.**
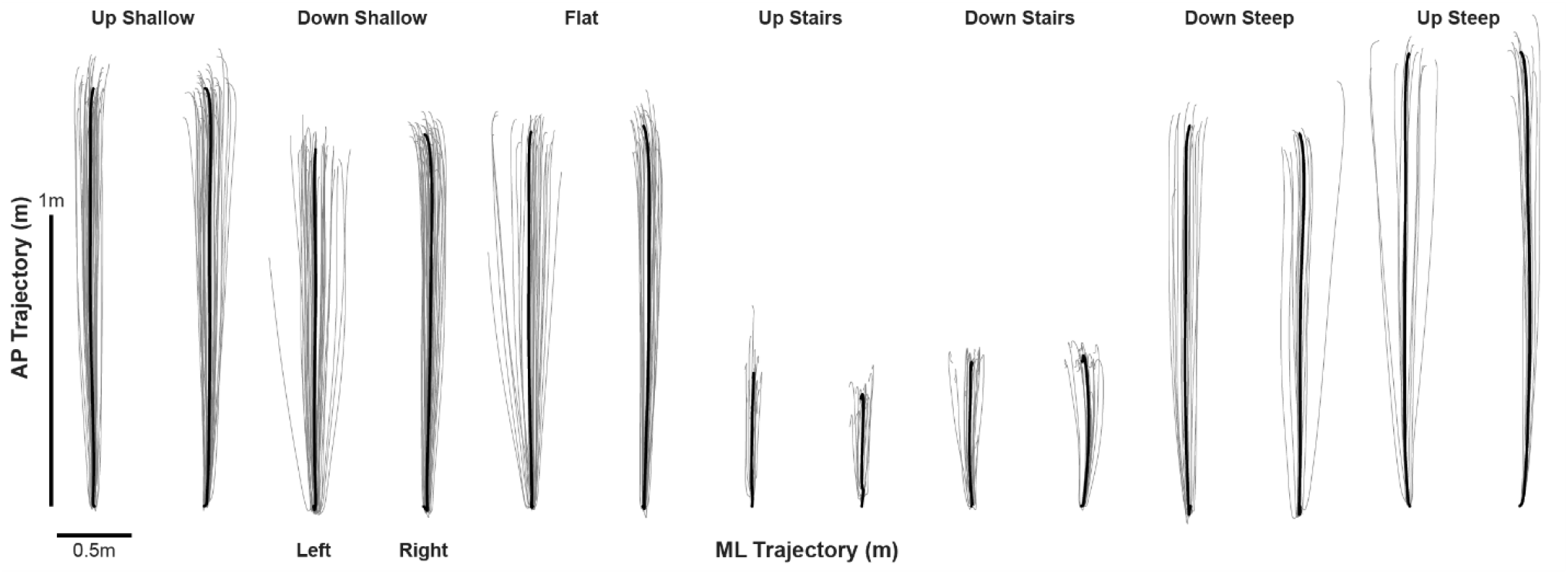
Left and right foot trajectory data from a representative participant during each terrain condition. Light grey illustrates individual stride trajectories while the thick black line represents the ensemble average of the trajectories.

### Statistical analysis

Trajectories were examined visually and presented as waveforms across each condition. The primary results were the discrete variables which we extracted as the mean and standard deviation of all the strides taken within each condition. The statistical tests comparing conditions are considered a secondary analysis given the hypothesis generating goals of this study.

To examine statistical differences in the discrete variables we performed one-way repeated measures analyses of variance, with the repeated effect being the walking condition. Non-sphericity was addressed with a Greenhouse-Geiser correction. The effect size was evaluated using the generalized eta square 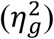. A significant main effect for condition (a = 0.05) resulted in pairwise paired t-tests corrected for multiplicity using a Hommel correction (30). The limb side (right or left) was not a significant effect in any model except sagittal plane foot angle at heel strike; therefore, models were run without the limb factor and with data combined from both limb sides. If the residuals of a given test were non-normal based on QQ-plots and a Shapiro-Wilks test, non-parametric tests were used (Friedman’s ANOVA and Wilcoxon Signed Rank Test). All statistical analyses were performed using R-Stats (v 4.3).

#### Results

Ten healthy adults enrolled in the study with mean (standard deviation) age of 25.9 years (2.0), height of 1.73m (6.3) and body mass of 72.3kg (12.9). Five participants were female and five were male and all indicated their right limb as dominant.

Foot trajectories largely followed a similar shape for slope and flat walking conditions with the stair conditions being much shorter (Fig 4). Modest differences were observed between the right and left limb with regard to the shape of foot trajectory, mainly in the mediolateral components. Discrete variables are reported in Table 2, while full statistical results are included in the Supplementary Tables.

**Table 2.** Discrete biomechanical outcomes as mean (95% Confidence Interval) for each walking condition.

| **Outcome** | **All Conditions** | **Descend Shallow** | **Descend Steep** | **Descend Stairs** | **Flat ground** | **Ascend Shallow** | **Ascend Steep** | **Ascend Stairs** |
| --- | --- | --- | --- | --- | --- | --- | --- | --- |
| **Stance time (s)** | 0.66 (0.64, 0.68) | 0.64 (0.62, 0.66) | 0.61 (0.59, 0.64) | 0.69 (0.62, 0.77) | 0.66 (0.64, 0.69) | 0.66 (0.63, 0.68) | 0.66 (0.63, 0.69) | 0.70 (0.64, 0.77) |
| **Stance time SD (s)** | 0.05 (0.04, 0.06) | 0.04 (0.03, 0.06) | 0.04 (0.03, 0.04) | 0.09 (0.06, 0.12) | 0.04 (0.03, 0.05) | 0.04 (0.04, 0.05) | 0.05 (0.03, 0.08) | 0.04 (0.04, 0.05) |
| **Swing time (s)** | 0.39 (0.37, 0.40) | 0.40 (0.38, 0.41) | 0.38 (0.37, 0.40) | 0.28 (0.25, 0.30) | 0.41 (0.39, 0.43) | 0.41 (0.40, 0.43) | 0.42 (0.41, 0.44) | 0.40 (0.37, 0.42) |
| **Swing time SD (s)** | 0.03 (0.03, 0.04) | 0.04 (0.03, 0.05) | 0.03 (0.02, 0.03) | 0.04 (0.03, 0.05) | 0.03 (0.02, 0.04) | 0.04 (0.03, 0.04) | 0.04 (0.02, 0.05) | 0.03 (0.02, 0.04) |
| **Cadence (strides/min)** | 61.45 (59.85, 63.05) | 58.88 (57.54, 60.21) | 63.61 (61.84, 65.37) | 70.37 (64.04, 76.69) | 57.84 (55.84, 59.85) | 57.63 (55.79, 59.47) | 58.58 (56.70, 60.47) | 63.24 (58.13, 68.35) |
| **Stride length (m)** | 1.06 (0.99, 1.14) | 1.17 (1.10, 1.23) | 1.18 (1.12, 1.25) | 0.59 (0.56, 0.61) | 1.23 (1.17, 1.29) | 1.33 (1.27, 1.38) | 1.39 (1.33, 1.45) | 0.56 (0.54, 0.58) |
| **Stride length SD (m)** | 0.11 (0.10, 0.12) | 0.12 (0.09, 0.15) | 0.09 (0.07, 0.10) | 0.08 (0.05, 0.10) | 0.10 (0.07, 0.13) | 0.13 (0.10, 0.15) | 0.15 (0.10, 0.21) | 0.12 (0.11, 0.14) |
| **Stride width (m)** | 0.06 (0.06, 0.07) | 0.08 (0.06, 0.09) | 0.07 (0.06, 0.09) | 0.06 (0.05, 0.06) | 0.07 (0.06, 0.07) | 0.06 (0.06, 0.07) | 0.07 (0.06, 0.07) | 0.05 (0.04, 0.06) |
| **Stride width SD (m)** | 0.05 (0.05, 0.06) | 0.06 (0.05, 0.07) | 0.06 (0.05, 0.07) | 0.05 (0.04, 0.05) | 0.06 (0.05, 0.06) | 0.05 (0.05, 0.06) | 0.05 (0.04, 0.06) | 0.04 (0.03, 0.05) |
| **Sagittal plane foot angle at IC (°)** | 13.67 (10.09, 17.25) | 25.36 (23.39, 27.33) | 26.88 (25.18, 28.58) | -15.20 (-16.72, -13.68) | 23.58 (21.80, 25.36) | 21.28 (19.41, 23.14) | 16.32 (15.00, 17.63) | -2.53 (-3.80, -1.26) |
| **Sagittal plane foot angle SD at IC (°)** | 5.59 (5.22, 5.96) | 5.05 (4.67, 5.43) | 7.08 (6.56, 7.60) | 7.29 (6.62, 7.95) | 5.15 (4.68, 5.63) | 5.16 (4.65, 5.66) | 6.21 (5.38, 7.04) | 3.18 (2.81, 3.56) |
| **Horizontal plane foot angle at midstance (°)** | -11.43 (-12.80, -10.05) | -12.74 (-17.01, -8.48) | -11.48 (-14.75, -8.22) | -11.88 (-16.48, -7.28) | -12.97 (-16.81, -9.13) | -10.75 (-13.51, -7.99) | -9.63 (-12.83, -6.43) | -10.52 (-14.42, -6.62) |
| **Horizontal plane foot angle SD at midstance (°)** | 6.14 (5.30, 6.98) | 5.74 (3.80, 7.69) | 7.21 (5.99, 8.43) | 9.97 (7.37, 12.57) | 4.31 (2.18, 6.44) | 4.27 (2.70, 5.84) | 3.97 (2.32, 5.63) | 7.49 (5.62, 9.35) |

Abbreviations: SD, standard deviation; IC; initial contact. Positive sagittal plane foot angles indicate the toe was above the heel while positive horizontal plane foot angles indicate the toe is more medial than the heel (toe-in).

**Figure 4.**
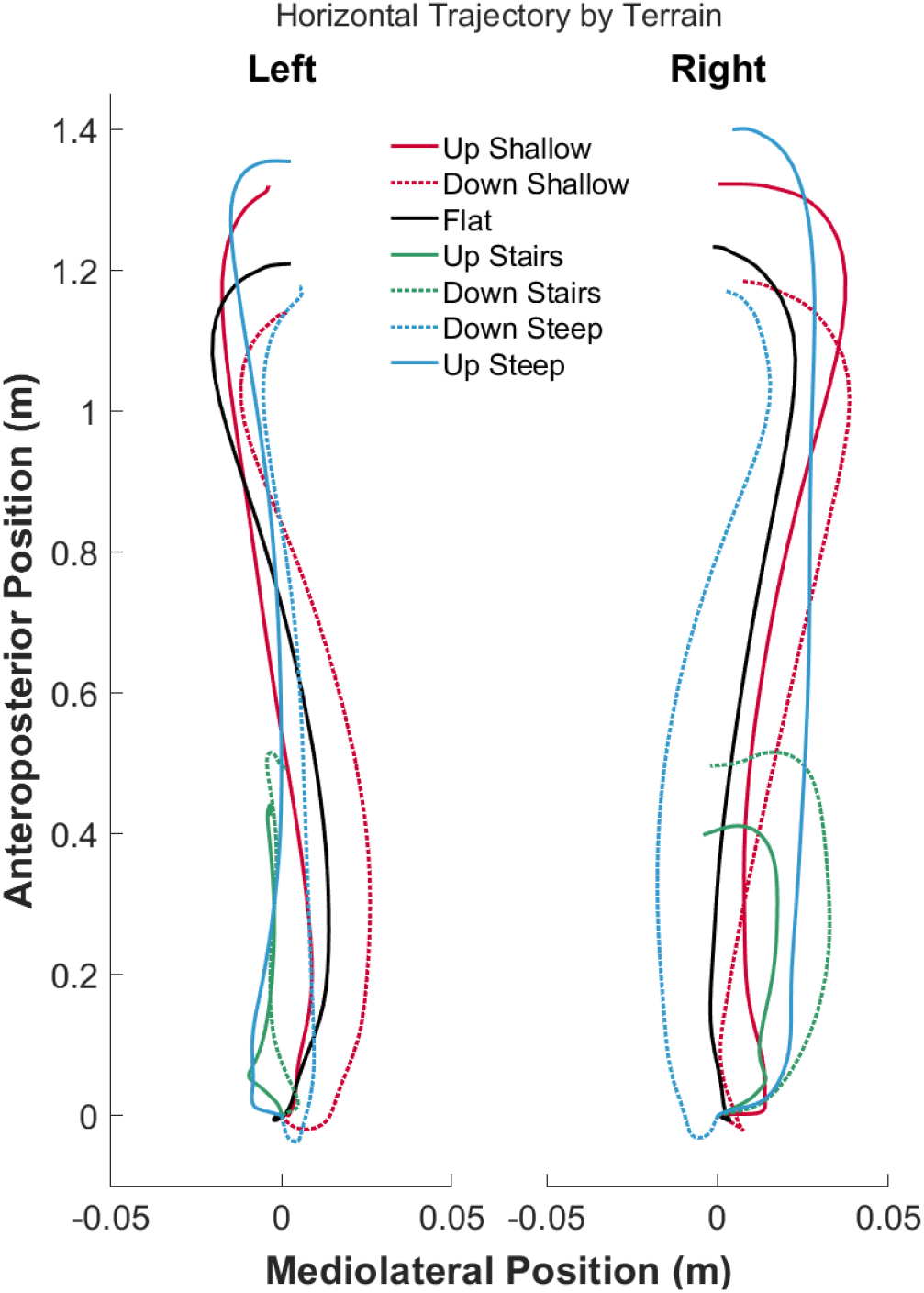
Mean foot trajectories across all participants and conditions in the mediolateral and anteroposterior directions. Solid lines are flat or ascending conditions, while dashed lines are descending conditions.

Several gait event timing outcomes significantly differed across the terrain conditions. Swing time (*F*=55.96, *df*=2.40/21.58, *p*<0.001, 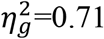) was shortest while descending stairs and steep slopes (*p*<0.006) and the variability (*Q*=13.0, *p*=0.043, *W*=0.217) significantly varied relative to the grand mean but no pairwise comparisons were significant (*p*>0.097). We did not detect differences in stance time (*F*=2.98, *df*=1.36/12.65,*p*=0.102, 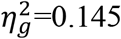) or stance time variability (*Q*=12.26, *p*=0.056, *W*=0.204). Cadence was significantly different across conditions (*Q*=40.157, *p*<0.001, *W*=0.669) with descending stairs and steep slopes requiring the highest cadence (*p*≤0.025). A single participant was a large outlier in all their spatiotemporal outcomes, so we refit the models with this participant data removed (see Supplementary Files). While the results were largely the same, the omnibus test for stance time was now significant (*Q*=23.57, *p*<0.001, *W*=0.44) though after corrections for multiplicity, only one significant comparison was detected with descending steep slopes relative to flat ground (*p*=0.008).

Both stride length (*Q*=55.33, *p*<0.001, *W*=0.922) and width (*Q*=25.67, *p*<0.001, *W*=0.43) were significantly different across conditions. Expectedly, stride length was shortest during both stair conditions given the physical dimensions (comparisons all *p*<0.001). Every other comparison was significant (*p*≤0.004), though ascending and descending stairs resulted in similar stride lengths (*p*=0.222). Stride width was generally smallest during the stair conditions (*p*<0.044) compared to the others, particularly for ascending stairs. The standard deviation of stride length (*Q*=12.99, *p*=0.043, *W*=0.22) was highest while ascending stairs and slopes compared to the descending condition, but the only significant comparison was between ascending stairs and descending steep slopes (*p=*0.005). Meanwhile, stride width standard deviations (*F*=4.40, *df*=6/54, *p*=0.001, 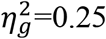) differed from the grand mean but we could not detect significant pairwise comparisons (*p*>0.062).

Different sagittal plane foot angles were used in all conditions (*Q*=57.47, *p*<0.001, *W*=0.96). When navigating stairs, participants contacted the ground with their forefoot first (negative angle), while the other conditions required a ‘heel first’ strategy depicted by positive foot angles (*p*<0.03 for all comparisons). The steeper the slope, the smaller the angle when ascending with the opposite effect for descending slopes (comparisons all *p*<0.001). The standard deviation of sagittal plane foot angles were different among conditions as well (*Q*=48.42, *p*<0.001, *W*=0.81) with ascending stairs exhibiting the lowest variability across all conditions (*p*<0.008) and descending stairs and steep slopes exhibiting the highest (*p*<0.014). Lastly, horizontal plane foot angle during midstance were significantly different based on the omnibus test (*Q*=15.0, *p*=0.021, *W*=0.25), but no significant pairwise comparisons were detected (*p*>0.067), except that ascending steep slopes required a significantly smaller angle than flat walking (*p*=0.008). Overall, participants used an externally rotated foot angle during each of the walking conditions. The standard deviation of horizontal plane foot angle varied across conditions (*F*=6.04, *df*=2.63/23.68, *p*=0.004, 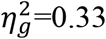), though the only significant comparisons showed that ascending shallow slopes had lower variability than descending slopes or stairs (*p*<0.014).

## Discussion

The increasing use of wearable IMU systems to capture gait in the real-world necessitates a better understanding of how locomotion is controlled and expressed in the varied terrain that we often encounter. Our study evaluated how healthy adults adjust their foot biomechanics while walking on stairs, flat ground, and sloped terrain. In order to collect data while walking on these real-world terrain conditions, we used a custom IMU embedded in the sole of the shoes, though our results would apply to any shoe-mounted IMU system with appropriate calibration procedures. The primary contribution of this study is the direct comparison of foot trajectory, orientation, and spatiotemporal metrics across seven different terrain conditions. Many of the outcomes we investigated were different between these conditions, with the most notable differences arising from stair navigation. These results and the extant literature demonstrate that different terrain generate unique foot biomechanics during locomotion, which should be accounted for when analyzing real-world data, particularly when there is limited knowledge of what terrain someone was walking on.

Adjusting stride length while navigating different terrain is not unusual. Our results support this given that nearly every condition exhibited different stride lengths, highlighting the sensitivity of this measurement to terrain geometry. In an observational study of pedestrians offloading a ferry, people descending a ramp showed a clear reduction in stride length compared to those ascending the same ramp (31). The authors hypothesized that this arose from different frictional demands of the foot-ground interface, where a slip is more likely during descent. Adapting stride length to avoid a slip during downhill walking is logical, but a lengthened stride in other situations could come at a cost. Previous research using both modelling and human experimental results during level ground walking showed that mechanical work and metabolic costs increased by the fourth power relative to a unit increase in step length (32), while other work reported a 3-fold increase in knee joint power during uphill walking at longer stride lengths (33). That is, adopting a longer stride length could require greater energy (34). Despite this, our participants opted to increase stride length by approximately 8-14% while ascending slopes compared to level ground. This stride length increase appears to be concomitant with an increase in forward trunk and pelvis inclination and is thought to be a strategy that can aid in whole body forward momentum (35). From our results, stride lengths are readily adapted to different terrain conditions, though some may result in higher metabolic costs. It would be important to categorize terrain conditions when comparing large real-world datasets as there are likely differential changes in stride length simply due to terrain.

As lateral balance needs to be actively controlled while walking (7, 36), adjusting step width to accommodate different terrain could be expected. However, we did not have the ability to measure step width; an estimation of balance control with relevance to functional status or aging (37, 38). Instead, we could measure stride width, which quantifies the mean change in mediolateral position of a single foot from stride to stride. Generally, we did not detect significant differences in the mean stride width across conditions after our *p*-value correction, though stair navigation had the narrowest widths by nearly 11-34% and the lowest variability. The smaller width changes from stride to stride while navigating stairs may reflect a strategy of more constrained motor control due to a more cognitively demanding task (39) or instability, compared to the other walking conditions.

Quantifying the orientation of the foot during gait has several clinically relevant implications. The sagittal foot angle just before contact will dictate how the foot will impact the ground, which could provide information on the potential for a slip incident (40). In the controlled settings of a laboratory, measurements of sagittal plane foot angle at initial contact suggest people use separate strategies when placing their foot on flat ground and slopes compared to stairs. Flat and sloped walking are usually associated with a heel-first contact with mean angles ranging from 10-32° (40-42). Our data largely corroborates these past findings, while also showing that the angle increased as the ascending slope angle increased, and the opposite occurring during descending.

Navigating stairs required different foot biomechanics when compared to level ground walking. While not the only strategy, it is often observed that the toe contacts the ground first (toe-below heel) both when ascending (-4.7°) and descending (-16.6°) (43). Although our specific foot angle values differed (-2.5° and -15.2°, respectively in our study) compared to Riener, Rabuffetti (43), we did observe the same trend relative to flat and sloped ground. Contacting toe-first during descending stairs is likely a strategy to dissipate the vertical forces during stance but may also arise due to the horizontal length of the individual stair relative to the foot length. The physical constraints of stairs also demand high knee and hip flexion while ascending, placing too great a demand on dorsiflexion range of motion to achieve a heel-first contact (43). Apart from different mean angles, we also observed lower sagittal foot angle standard deviation during stair ascent. This could represent the maintenance of more control over foot trajectory and foot-ground contact to minimize the risk of catching the toe on a stair during the swing phase and inducing a trip. Lastly, a possible source of between-subject variability in sagittal foot angle measures during our ascent conditions more broadly could be dorsiflexion range of motion. Lower inherent range of motion could limit sagittal plane foot angles relative to the ground (44). It is unlikely this played a prominent role in our slope conditions given we used a slope of 6° (Kwon and Shin (44) = 20°), though the stairs were 36° which may have interacted with individual dorsiflexion range of motion. Future studies using steeper inclinations than we used may need to consider this when stratifying a sample population or excluding participants in order to avoid unwanted variance in their results.

The horizontal plane foot angle during midstance was similar throughout the study. The magnitude of horizontal plane foot angles can vary quite widely among individuals, partly due to boney anatomy (45). We observed a range from -9.6° to -13 on average indicating an externally rotated foot relative to the direction of walking, which was well within usual measurements (46-48). Tracking this metric using wearable IMUs has utility in gait modification interventions where people learn to increase or decrease the angle relative to the direction of walking (usually called foot progression angle) (49, 50). In recent years, our group has implemented wearable technology to monitor this metric in real-world settings (26), where the present studies results will be valuable in contextualizing the data collected while participants walked on different terrain in their community, over weeks and months.

The timing of gait is closely related to the speed and cadence (51, 52), which is liable to change while navigating different terrain in the real world. The most notable difference was a significantly shorter time spent in swing when descending steep slopes and stairs, concomitant with a higher cadence. This likely arose from the terrain lending to faster overall speeds and shorter stride length. Additionally, we observed a larger ratio of stance to swing times (longer stance relative to swing) while participants navigated stairs, which was similar to past research (43, 53). However, this was opposite of the findings from Nadeau, McFadyen (54); perhaps due to the fact our stair condition had more stairs (4 vs 16) and were slightly steeper (33° vs 36°), which may have prompted participants to adopt a different strategy.

Several limitations need to be considered when interpreting our results. We examined a sample of healthy adults, all of which had no sensorimotor deficits or neuromuscular conditions that could affect their ability to walk. This allowed for a controlled evaluation of the effects that terrain induces on gait. However, we could not capture the potential terrain-dependent effects that could arise in other populations, such as older adults. For example, older adults tend to reduce stride lengths with increasing slopes or multi-surface terrain compared to healthy adults (55, 56). We only collected data on slopes of shallow (3°) and steep (6°) inclination. There are situations in our built and natural world that may require navigating slopes of much steeper conditions, or perhaps slopes in between the conditions we examined. Ultimately, we used representative and publicly accessible terrain conditions but a more refined comparison with smaller increments of slope inclination could warrant comparisons with more resolution. We did not explicitly control walking speed during our data collections. Instead, we opted to collect data across a range of speeds governed by the participants’ own preferences in each condition. This increased the between-subject variability in our data but captured the natural differences in how people navigate different terrain at different speeds. It is likely that a more strict control of speeds could result in more detectable differences between these terrain conditions, as can be found in laboratory experiments (54). Our estimates of foot orientation rely on accurate identification of the midstance event, for which we opted to use separate gait event detection algorithms for stair navigation and slopes or flat walking. It is assumed that foot acceleration is near zero at this timepoint, which may not always occur during stair navigation or faster walking. An alternative method of resetting orientation at each midstance may be required to improve the accuracy of estimates while walking on stairs or at higher velocities. Finally, while we conducted our study out of the laboratory, it was still reasonably controlled as participants walked on linear paths and in contexts with limited distraction. Continued investigations in this area should also consider the context of a walking bout (e.g., commute, walking the dog, shopping etc.), surrounding environment (e.g., busy side walk, park path) and evaluate more realistic scenarios as the context of a walking bout may influence gait dynamics (57).

Taken together, our results indicate that it is important to consider terrain when collecting gait data in ecologically valid environments or in unsupervised conditions. As wearable IMUs become the dominant data collection modality in gait research, the opportunities for real-world collection continue to expand. Given the inherent and uncontrollable variability in real-world, unsupervised conditions, it is important to understand the extent to which different data collection conditions could impact results. Our study highlights these often small, but potentially meaningful differences based on the data we collected on a set of common real-world terrain.

